# Linking nitrogen load to the structure and function of wetland soil and rhizosphere microbial communities

**DOI:** 10.1101/197855

**Authors:** Eric R Hester, Sarah F. Harpenslager, Josepha MH van Diggelen, Leon L Lamers, Mike SM Jetten, Claudia Lüke, Sebastian Lücker, Cornelia U Welte

**Affiliations:** Department of Microbiology, Radboud University, Nijmegen, The Netherlands; Department of Aquatic Ecology and Environmental Biology, Radboud University, Nijmegen, The Netherlands; School of Biological and Chemical Sciences, Queen Mary University, London, United Kingdom; B-WARE Research Centre, Nijmegen,The Netherlands

**Keywords:** greenhouse gas, microbial community function, Opitutales, Acidobacteria, Sphingobacteriales, wetlands, *Juncus acutiflorus*, nitrogen

## Abstract

Wetland ecosystems are important reservoirs of biodiversity and significantly contribute to emissions of the greenhouse gases CO_2_, N_2_O and CH_4_. High anthropogenic nitrogen (N) inputs from agriculture and fossil fuel combustion have been recognized as a severe threat to biodiversity and ecosystem functioning such as control of greenhouse gas emissions. Therefore it is important to understand how increased N input into pristine wetlands affects the composition and activity of micro-organisms, especially in interaction with dominant wetland plants. In a series of incubations analyzed over 90 days, we disentangle the effects of N fertilization on the microbial community in bulk soil and the rhizosphere of *Juncus acutiflorus*, a common and abundant graminoid wetland plant. We observed an increase in greenhouse gas emissions when N is increased in incubations with *J. acutiflorus*, changing the system from a greenhouse gas sink to a source. Using 16S rRNA amplicon sequencing and metagenomics, we determined that the bacterial orders Opitutales, Subgroup-6 Acidobacteria and Sphingobacteriales significantly responded to high N availability and we hypothesize that these groups are contributing to the increased greenhouse gas emissions. These results indicated that increased N input leads to shifts in microbial activity within the rhizosphere, severely altering N cycling dynamics. Our study provides a framework for connecting environmental conditions of wetland bulk and rhizosphere soil to the structure and metabolic output of microbial communities.

## Introduction

Wetlands are globally impacted by agricultural industry through the leaching of various nitrogen (N) forms such as nitrate (NO_3_^−^), and by increased N deposition as a result of high N emissions from fossil fuel burning and agriculture [Galloway *et al.*, 2008]. Furthermore, due to reduced oxidation under stagnant, waterlogged conditions, these systems show increased availability of ammonium (NH_4_^+^) (Britto and Kronzucker, 2002). The strongly increased anthropogenic N input influences ecosystem degradation by contributing to biodiversity loss and altering (mostly increasing) greenhouse gas fluxes such as nitrous oxide (N_2_O), methane (CH_4_) and carbon dioxide (CO_2_) (Bobbink *et al.*, 1998; Liu and Greaver, 2009]; Van den Heuvel *et al.*, 2011; Soons *et al.*, 2016).

The abundance, composition and activity of micro-organisms strongly influence the biogeochemical cycling of wetland nutrients, particularly those resulting in emissions of greenhouse gases (Lamers *et al.*, 2012; Philippot *et al.*, 2009). Specifically, N_2_O emission may increase due to lowering of pH affecting the activity of incomplete denitrifiers (Brenzinger *et al.*, 2015; Van den Heuvel *et al.*, 2011; Liu and Greaver, 2009). CH_4_ emissions can increase due to competitive inhibition of the key enzyme of aerobic methanotrophs, methane monooxygenase (MMO), by elevated NH_4_^+^, osmotic stress of methanotrophs, or through the stimulation of methanogenic archaea (King and Schnell, 1998; Bodelier and Laanbroek, 2004; Dunfield and Knowles, 1995). Finally, the rate of soil C loss can increase as a result of N addition through the stimulation of heterotrophic respiration (Bragazza *et al.*, 2006). Although it is well established that microbial processes are important drivers of ecosystem functions, such as controls on greenhouse gas emissions and nutrient cycling, there is a lack of understanding of how these functions are linked, both to the environmental conditions and to the composition of the microbial community (Philippot *et al.*, 2009).

Wetland plant roots influence the soil region surrounding the root, known as the rhizosphere, by altering the availability of oxygen, organic matter, and organic plant exudates (Smith and Delaune, 1984; Abou Seada and Ottow, 1985; Bardgett and van der Putten, 2014). The total area of soil influenced by roots can be considerable, meaning that this definition of the rhizosphere may extend to the vast majority of the upper soil layer (Robinson et al., 2003). The rhizosphere is an active, complex ecosystem where viruses, bacteria, archaea, fungi and protozoa interact with plant roots (Fierer et al., 2007b). These microorganisms significantly contribute to nutrient cycling and ecosystem structure by channeling energy into higher trophic levels (reviewed in Curl & Harper 1990; Hinsinger et al. 2009).

While the rhizosphere has been studied for decades, the effects of eutrophication on the plant-microbe interactions are of more recent interest. Specifically, it is of interest how N availability influence plant physiology and ultimately C and N cycling in the rhizosphere. On the global scale, soil microbial communities differ depending on the regional and local N regime; although, the diversity of these communities does not seem to vary much (Fierer *et al.*, 2012). Interestingly, variation in microbial community composition seems to be predictable based on local nutrient regimes (Leff *et al.*, 2015; Ramirez *et al.*, 2012). Even though these studies demonstrate the link between nutrient loading and community structure, they do not demonstrate how changes in the microbial community are functionally relevant to the ecosystem.

To build dynamic models of plant-microbe interactions, it is necessary to gain a robust understanding of the connection between environmental conditions (i.e., N availability) and microbial community structure and function (i.e., the bulk biological processes resulting in greenhouse gas emissions). In this study, we aimed at assessing the impact of increased N input into wetland systems on therhizosphere microbial community and its functions related to greenhouse gas production. To achieve this, we used *Juncus acutiflorus* (Sharp-flowered Rush), a very common graminoid plant in European wetlands that forms a dense vegetation and is known for radial oxygen loss from roots (ROL; Lamers et al. 2012). Furthermore, it has a high tolerance for increased N inputs (van Diggelen *et al.*, 2016).In a longitudinal study we determined greenhouse gas emissions increase as a result of N addition in incubations with *J. acutiflorus*, but not in incubations with only bulk wetland soil, under controlled stable experimental conditions. Additionally, functional responses were linked to shifts in the dominant members of the microbial community. We hypothesize that certain key microbial groups contribute to greenhouse gas emissions, either directly or indirectly through the food web. Our study takes the first steps toward a predictive understanding of microbial dynamics within the rhizosphere, linking nutrient load, microbial community structure and function.

## Materials and Methods

### Sample Collection and experimental set up

Plants and sandy soil were sampled from the Ravenvennen (51.4399 N, 6.1961 E) in Limburg, The Netherlands (August, 2015) and returned to the Radboud University greenhouse facilities for conditioning. The Ravenvennen is a protected marshy area consisting of sandy soil, rich in vegetation with a high prevalence of *Juncus spp*. Plants were removed from soil, rhizomes were cut into eight 2 cm fragments and reconditioned on hydroculture in a nutrient rich medium (as described in Hoagland & Arnon 1950). After sufficient root development (to approximately 25 cm after 2 weeks), eight plants and eight bulk soil incubations were randomly assigned to high or low nitrogen experimental groups (Supplementary table 1; Supplementary figure 1). Soil collected from the field, was homogenized and sieved to remove any contaminating roots and potted. The reconditioned plants were transferred to pots with a diameter of 19 cm at the base, 26 cm at the top and a height of 19 cm containing the prepared soil, moved to an indoor water bath set to 15 ° C (cryostat, NESLAB, Thermoflex 1400, Breda, The Netherlands) and cultivated with a day/night cycle of 16 hours light and 8 hours dark (Master Son-T PiaPlus, Philips, Eindhoven, The Netherlands). Pots were kept waterlogged with a 2 cm water layer on top. A drip-percolation based system ensured a constant supply of nutrients. The low N input nutrient solution contained 12.5 μM NH_4_NO_3_, corresponding to an N loading rate of 40 kg N ha^−1^ yr^−1^. The high N input solution contained 250 μM NH_4_NO_3_, corresponding to 800 kg N ha^−1^ yr^−1^. These rates fall within N loading of wetlands in agricultural catchments, thus represent contrasting extremes (Verhoeven et al., 2006).

### Incubation measurements

Five representative *J. acutiflorus* specimens were harvested for initial measurements of plant dry weight, C:N ratios. At the final time point (T_f_ = 90 days), all plants were harvested to measure dry weight and C:N ratios of roots, shoots and rhizomes. Pore water was extracted using 0.15 μm porous soil moisture samplers (SMS rhizons, Rhizosphere Research Products, Wageningen, The Netherlands) and measured over the course of the experiment to determine inorganic nutrients as well as metals using an Autoanalyzer (Autoanalyzer 3, Bran+Luebbe, Germany) and ICP-OES (iCAP6000, Thermo Scientific,Waltham, MA). To reduce the impact of soil heterogeneity, samples were extracted in duplicate and mean values were calculated.

### Greenhouse gas measurements

To determine greenhouse gas fluxes, a cylindrical transparent collection chamber (7.5 x 30cm) was used to measure accumulation or depletion of CO_2_, CH_4_ and N_2_O in the headspace. CO_2_ and CH_4_ fluxes were measured at T_m_ (45 days) and T_f_ and N_2_O fluxes were measured at T_f_. Fluxes were measured using a Picarro G2308 NIRS-CRD greenhouse gas analyzer (Picarro Inc., Santa Clara, CA, USA). Fluxes were determined by fitting a smoothed spline to the time series using the R function *sm.spline* from the pspline package and the average rate of change was calculated (Ramsay *et al.*, 1997).

### Denitrification potential

To measure denitrification potential, two soil slurries were made from each experimental pot by mixing 50g soil with 100mL milliQ water, divided into control and experimental bottles and made anoxic by flushing with argon gas. Bottles were pre-incubated overnight at 15°C to allow for residual unlabeled NO_3_^−^ to be consumed. A ^15^N-labeled NaNO_3_ solution was added to the experimental bottles to a final concentration of 500 μM and a KCl solution was added to the control bottles to a final concentration of 500 μM. Production of N_2_O and N_2_ were measured by taking samples 2, 7 and 22 h after adding substrate on a GC-MS (5975C, Agilent Technologies, Santa Clara, USA).

### DNA extraction, 16S rRNA Amplicon and Metagenomic sequencing

Soil was collected from three time points, one initial soil sample from the site, and T_m_ and T_f_ samples from each of the 16 incubations. A single core per pot was taken using a 1x7cm corer. DNA was extracted using the PowerSoil DNA Isolation Kit (MoBio, Carlsbad, CA, U.S.A.). 16S rRNA genes were amplified in triplicate reactions using IonTorrent sequencing adapter-barcoded primers 341F (CCATCTCATCCCTGCGTGTCTCCGACTCAGxxxxxxxxxxGATCCTACGGGNGGCWGCAG) and 785R (CCACTACGCCTCCGCTTTCCTCTCTATGGGCAGTCGGTGATGACTACHVGGGTATCTAA TCC) and pooled. The pooled amplicons were cleaned with Ampure beads (Beckman Coulter Inc., Fullerton, USA) and subsequently prepared for sequencing on the IonTorrent PGM using the manufacturer’s instructions (Life Technologies, Inc., Carlsbad, CA, USA).

From the same DNA samples, total DNA was sheared into approximately 400 bp fragments via sonication. Resulting fragments were prepared for sequencing following the manufacturer’s instructions with the Ion Plus Fragment Library Kit (Life technologies, Carlsbad, CA). Raw reads were submitted to NCBI and archived under the SRA accession number SRP099838.

### Data analysis

16S rRNA gene amplicons were quality filtered using QIIME v1.9 (Caporaso *et al.*, 2010). Quality controlled reads were then clustered into OTUs at a 97% identify and phylogenetically classified by utilizing the NINJA-OPS v1.3 pipeline (Al-Ghalith *et al.*, 2016). The reference database used for taxonomic assignment was the SILVA database version 123 (Quast *et al.*, 2013). The resulting OTU table was used for downstream analysis in R (R Core Team, 2016). Count data was normalized to relative abundances to account for differing sequence depth between samples and a square root transformation was applied. The *vegan* R package was used to calculate Shannon diversity with the *diversity* function, Bray-Curtis dissimilarity matrices with the *vegdist* function, and to estimate compositional variance with the *betadisper* function (Oksanen *et al.*, 2015). Principal component analysis (PCA) was performed using the *princomp* function in R. The *RandomForest* R package was used for classification and regression (Liaw and Wiener, 2002). Linear models were fit with the *glm* function in the *stats* package. Metagenomic reads were quality filtered (Q > 25) and small fragments (< 100bp) were removed using PrinSeq (Schmieder and Edwards, 2011).

The metagenomic reads were compared to custom nitrogen and methane cycling protein databases and the NCBI nr databases with Diamond (Buchfink *et al.*, 2014; Lüke *et al.*, 2016). A bit score ratio (BSR) between the hit to the custom databases and to the NCBI nr database was used to identify false positives hits. A strict BSR of 0.85 was used as a cutoff. Gene abundances were normalized and expressed relative to the single copy RNA polymerase *rpoB* gene abundance. These relative values were then scaled for comparison within genes. Reads from all metagenomes were assembled using metaSPAdes (version 3.7; Bankevich et al. 2012) and resulting contigs were compared against all publicly available Bacteroidetes, Acidobacteria and Verrucomicrobia genomes in the NCBI database using Blastn. Furthermore, contigs were assessed for the presence of N or CH_4_ cycling genes by comparing them with Diamond to the previously mentioned N and CH_4_ cycling custom databases.

## Results

### Plant physiology

*J. acutiflorus* and bulk soil were incubated over a course of 90 days. The soil collected from the sampling site and used in the incubations was a sandy soil with low organic matter content. Soil samples were taken at an initial time point (T_0_), a mid-point (T_m_; t= 45 days) and final time point (T_f_; t= 90 days) (Supplementary Table 1). By T_m_, *J. acutiflorus* incubations had significant root development throughout the incubated soil, and as a result the rhizosphere was sufficiently sampled such that the soil sampled was clearly dominated by root biomass. To determine the N utilization of the plants and to identify growth responses to N inputs, the total dry weight biomass of roots, rhizomes and shoots and total N and C content of *J. acutiflorus* tissue were measured from plants at T_f_. Although there was no significant difference in total biomass and root:shoot ratio of *J. acutiflorus* between incubations, the average total N content of plant tissue was approximately twice as high (65 mg g^−1^) in incubations with a high N input (t = 2.66; p = 0.037; Supplementary Table 2). Correspondingly, total C:N (averaged across the whole plant) was significantly higher in *J. acutiflorus* incubations with a low N input (t = −2.964; p = 0.009; Supplementary Table 2). Interestingly, this elevated C:N ratio was observed only for rhizome and shoot tissue, while the root C:N did not significantly differ between incubations (Supplementary Table 2).

### Greenhouse gas fluxes

To link greenhouse gas fluxes with microbial community structure, gas flux measurements were performed at the same time points as soil sampling. Greenhouse gases were measured in both light and dark conditions, at T_m_ and T_f_ for CO_2_ and CH_4_, and at T_f_ for N_2_O (Figure 1). Bulk soils generally did not have significant greenhouse gas fluxes (fluxes were not significantly different from 0) and will not further be discussed here. In the *J. acutiflorus* incubations, CO2 fluxes followed a day-night rhythm. Daytime CO_2_ fluxes were generally negative, indicating net CO_2_ fixation, with the largest rates significantly higher in high N *J. acutiflorus* incubations at T_f_ (t = −5.28, p = 0.005; Figure 1A). Under dark conditions, CO_2_ fluxes were positive only under the high N treatment while other treatments were not significantly different from 0(t = 3.52, p = 0.01; Figure 1B). CH_4_ and N_2_O emissions did not vary between dark and light conditions and therefore these conditions will not be compared. CH_4_ fluxes increased from T_m_ to T_f_ and emissions tended to be highest in the *J. acutiflorus* incubations with a high N input, however there was large variability in this group (t = 2.165; p = 0.064; Figure 1C). N_2_O emissions were highest in the high N treatment (t = 2.56, p = 0.04; Figure 1D), while a negative N_2_O flux was observed in *J. acutiflorus* incubations receiving a low N input (Figure 1D).

**Figure 1.**
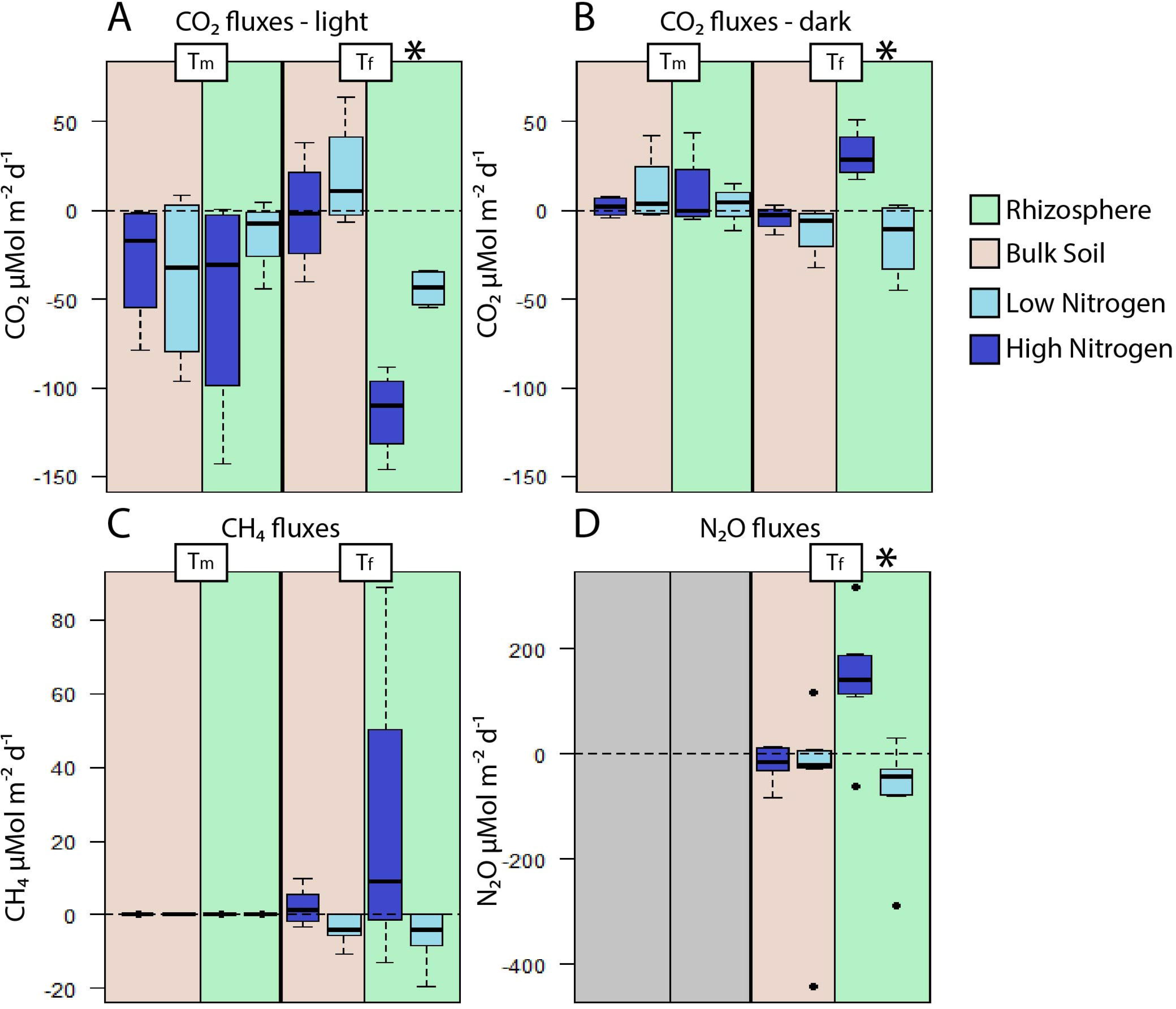
CO_2_, CH_4_ and N_2_O fluxes. Greenhouse gas fluxes were measured at a midpoint (T_m_) and final time point (T_f_) during the day incubation experiment. (A) CO_2_ light conditions, (B) CO_2_ dark conditions, (C) CH_4_ and (D) N_2_O. Asterisks denote significant differences (p < 0.05).

### Denitrification potential

To understand how increased N input influenced N cycling within bulk and *J. acutiflorus* rhizosphere soils, soil slurries were taken at T_f_ and their denitrification potential was measured. There was significantly higher N_2_O production from slurries originating from high N treatment soils (t = 2.41; p = 0.045; Supplementary Figure 2). There was no significant difference in the N_2_ production between high or low N treatments (t = 0.32; p = 0.75; Supplementary Figure 2). Additionally, the average N_2_:N_2_O ratio was approximately 10 times higher in low N input slurries (5.36 +/− 7.39; N_2_:N_2_O production) as compared to high N slurries (0.58 +/− 0.61), though not significantly different at p < 0.05 (t = −1.84; p = 0.11; Supplementary Figure 2).

### Microbial community structure

The v3-v4 fragment of the 16S rRNA gene was amplified and sequenced resulting in, on average, over 1100 post-quality control (QC) sequences per sample. Each sample contained on average 264 +/− 136 Operational Taxonomic Units (OTUs +/− s.d.). Rarefaction curves suggest that communities were sampled to capture the majority of the diversity (Supplementary Figure 3). Over the course of the incubation, the dominant microbial group changed (Figure 2A). Solibacteriales were most abundant at T_0_ and at T_m_, but by T_f_ Rhizobiales became the prominent group (Figure 2B). On average, microbial diversity increased between T_m_ and T_f_, (t = 2.516; p = 0.0176; Supplementary Figure 4A). Within each time point, diversity did not differ significantly between *J. acutiflorus* and bulk soil incubations, nor did N input have an impact (Supplementary Figure 4B+C). To assess how community composition varied across the different incubations, the Bray-Curtis dissimilarity index was used to calculate compositional differences between microbial communities. The compositional variation did not significantly vary between T_m_ and T_f_, indicating that community variability did not change within the different experimental groups across time (Supplementary Figure 5A-C). The most variable communities were observed for low N *J. acutiflorus* incubations at T_m_, which furthermore were significantly different from the low N input bulk soil incubations (Tukey’s HSD; p = 0.0184; Supplementary Figure 5B). At T_f_ there were no significant differences in community variation among bulk soil or *J. acutiflorus* incubations, or between low and high N loading. There were significant differences in overall community composition between high and low N treatment (PerMANOVA; p = 0.003), rhizosphere and bulk soil (p = 0.02), and midpoint and final time points (p < 0.001).

**Figure 2.**
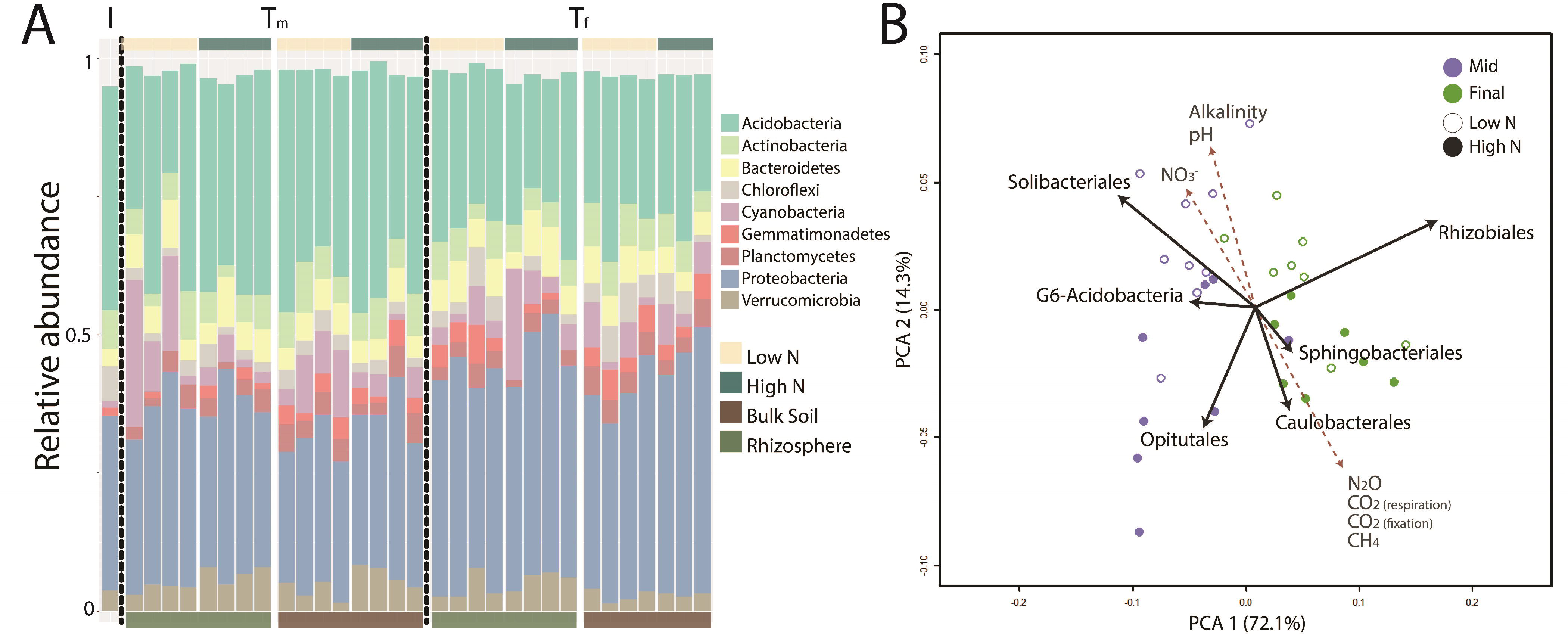
Microbial community structure and diversity.
(A) Overview of microbial community structure of the initial soil sample (I), *J. acutiflorus* rhizosphere and bulk soil incubations receiving high or low N input at midpoint (Tm) and final time point (T_f_). (B). Principal component analysis of the microbial community members distinguishing high and low N treatments and midpoint and final sampling time points. Points indicate individual samples taken. Red dashed arrows indicate environmental and gas fluxes that corresponded variation in microbial community member’s abundance along the respective axis.

### Linking microbial community members to function

In order to understand how the microbial community members were linked to environmental conditions and greenhouse gas emissions, a random forest classifier was used to identify microbial taxa whose abundance was affected by N input, time of sampling or presence of *J. acutiflorus.* Additionally, random forest was also used for regression to determine connections between abundance of these groups and environmental conditions or greenhouse gas fluxes, and these associations were further analyzed by fitting linear models.

The top three microbial groups that significantly responded to N input were the Opitutales (Verrucomicrobia) and Sphingobacteriales (Bacteroidetes), which were more abundant in the high N treatment group, and G6 Acidobacteria, which were more abundant in in the low N treatment (Figure 2B; Table 1). More specifically, the relative abundances of these three orders could be linked to N_2_O emissions (Table 1). Opitutales and Sphingobacteriales were positively associated with N_2_O fluxes, while a negative association was observed for the G6-Acidobacteria. In addition, Sphingobacteriales were correlated to CO_2_ fixation (Table 1).

**Table 1.** Correlations of microbial community members to environmental conditions and greenhouse gas fluxes. The mean relative abundance of top bacterial families distinguishing high v low N, rhizosphere v bulk soil or T_m_ v T_f_ sampling time points are indicated as is the t-test result and statistics. Additionally, the top environmental or functional traits correlated with these groups were reported along with linear model statistic

|  | t | p-value | Mean Relative Abundance |  | Correlate | adj R2 | coef | p-value |
| --- | --- | --- | --- | --- | --- | --- | --- | --- |
| *High versus low N* |  |  | *High N* | *Low N* |  |  |  |  |
| Opitutales | 4.17 | < 0.001 | 0.040 | 0.010 | N<sub>2</sub>O | 0.11 | 3.50E-04 | 0.012 |
| G6-Acidobacteria | -4.22 | < 0.001 | 0.007 | 0.020 | N<sub>2</sub>O | 0.19 | -3.18E-05 | 0.058 |
| Sphingobacteriales | 2.88 | 0.008 | 0.010 | 0.005 | N<sub>2</sub>O | 0.32 | 3.10E-05 | 0.016 |
|  |  |  |  |  | CO<sub>2</sub> (fixation) | 0.29 | 7.07E-05 | 0.011 |
| *Rhizosphere versus bulk* |  |  | *Rhizosphere* | *Bulk* |  |  |  |  |
| Caulobacterales | -3.46 | 0.002 | 0.052 | 0.032 | NO<sub>3</sub><sup>-</sup> | 0.21 | -8.50E-05 | 0.003 |
| *T<sub>m</sub> versus T<sub>f</sub>* |  |  | *T<sub>m</sub>* | *T<sub>f</sub>* |  |  |  |  |
| Rhizobiales | 6.66 | < 0.001 | 0.099 | 0.184 | CO<sub>2</sub> (respiration) | 0.27 | -6.40E-04 | 0.001 |
| Solibacterales | -4.76 | < 0.001 | 0.179 | 0.116 | Alkalinity | 0.26 | -2.00E-02 | 0.002 |

The top bacterial order distinguishing microbial communities from rhizosphere and bulk soil were the Alphaproteobacterial Caulobacterales, which were more abundant in the rhizosphere than in bulk soil and had a negative association with elevated NO_3_^−^ concentrations (Table 1). The Rhizobiales and Solibacterales orders of the Alphaproteobacteria class and Acidobacteria phylum, respectively, were most distinctive for the microbial communities sampled at T_m_ versus T_f_ (Figure 2; Table 1). Rhizobiales abundance was negatively associated with CO_2_ fluxes in dark conditions while the Solibacterales were correlated to pore water alkalinity, which is a proxy for anaerobic decomposition (Figure 2; Table 1).

### Soil metagenomics

In addition to the 16S rRNA gene, which cannot be linked to functional genes on their own, total DNA was sequenced from 5 soils with representatives from T_0_, and rhizosphere and bulk soil samples at T_m_ and T_f_ from the high N treatment. The goal of the metagenomic sampling was to survey the genetic potential of organisms that were most strongly influenced by N loading. In particular, we wanted to find support for the roles the taxa mentioned above have in the rhizosphere of *J. acutiflorus*. These libraries resulted in on average 1 million post-QC reads per library (Supplementary Table 3). Over 4.8 million soil metagenome reads were then assembled into 129,476 contigs with a maximum length of 23kbp and a mean length of 597bp (+/− 368bp). Assembled contigs were compared to publicly available bacterial genomes from the Bacteroidetes, Verrucomicrobia and Acidobacterial phyla to identify genome fragments derived from the species identified in our previous analysis. Across all metagenomes, 5454 reads mapped to 145 contigs which had high identity to a Subgroup 6 Acidobacterial genome (CP015136.1; 84.5 +/− 7.1% identity), 6831 and 22 reads mapped to 352 and 5 contigs which aligned to an Opitutales (CP016094.1; 85.5 +/− 7.9% identity) and Sphingobacteriales (CP003349.1; 86.3 +/− 7.7% identity) genomes respectively (Supplementary Table 4).

In order to survey genetic potential for N and C cycling in N amended samples, custom databases of genes involved in N and C cycling processes (Lüke et al., 2016) were used to identify metagenomic reads of major N *(amoA* and *hao*, involved in NH_4_^+^ oxidation; *narG, nirK, nirS, norB*and *nosZ*, involved in denitrification; *nrfA*, involved in dissimilatory nitrite reduction to ammonia; and *nifH*, involved in N fixation) and CH_4_ cycling genes (*pmoA and mmoX*, inovled in CH_4_ oxidation; *phnGHI* and *mcrA*, involved in methanogenesis), and their abundance in the high N incubations (abbreviations found in Supplementary Table 5). There were no *nirS* detected in the dataset and only two reads annotated as *mcrA* were detected in the metagenomes from *J. acutiflorus*. All other N and CH_4_ cycling genes were present.

## Discussion

Greenhouse gas emissions remain a global challenge. A mechanistic understanding of the factors that alter microbial community structure and function, such as increased N input, is important in developing management strategies for greenhouse gas emissions. This is particularly important in ecosystems as extensive as wetlands. With an estimated area of up to 12.8 million km^2^ worldwide, wetlands considerably contribute to the total terrestrial carbon storage (Zedler and Kercher, 2005; Nahlik and Fennessy, 2016). Here we studied the impact of increased N input on the microbial community and greenhouse gas fluxes from the rhizosphere of *Juncus acutiflorus*, a very common plant in European wetland ecosystems, and a model for other *Juncus* species globally. We found characteristic shifts in the microbial community structure and a stimulation of greenhouse gas fluxes in *J. acutiflorus* incubations in response to N input.

### Plant physiological shifts as a response to high N inputs

The plant plays a prominent role in the maintenance of the rhizosphere microbial community (Reinhold-Hurek *et al.*, 2015). Roots release oxygen through radial oxygen loss providing an oxic niche in otherwise anoxic wetland soils (Armstrong, 1971). Plants also release labile organic matter in the form of organic acids, neutral sugars and amino acids (Kamilova *et al.*, 2006; Jones, 1998).The composition of this organic matter structures the microbial community within the rhizosphere by providing different substrates for heterotrophic micro-organisms (Haichar *et al*, 2008). The exuded organic acids also acidify the surrounding soil, preventing many microbial species from thriving within the rhizosphere, but also modifying nutrient availability (Marschner *et al.*, 1987; Petersen and Böttger, 1991). The quantity of organic matter released is closely associated with photosynthesis rates. As plants are often N limited in natural systems, relieving this limitation promotes plant growth (Reich *et al*, 2006). In this study we observed that when incubated under high N input *J. acutiflorus* showed increased C fixation rates (Figure 1A) and plant tissue becomes saturated with N (Supplementary Table 2). This also suggests that *J. acutiflorus* without N limitation excretes larger amounts of labile carbon into the surrounding soil, which is also evident from the observed decreases in pore water pH in the high N incubations(Supplementary Figure 6). Additionally, due to root derived oxygen, increased nitrification rates could contribute to this observed drop in pH Lamers et al. 2012). Together, higher N input could result in higher photosynthetic rates in *J. acutiflorus* specimens, likely depositing larger amounts of organic matter into surrounding soil, stimulating the heterotrophic microbial community in return (Figure 2; Figure 3).

**Figure 3.**
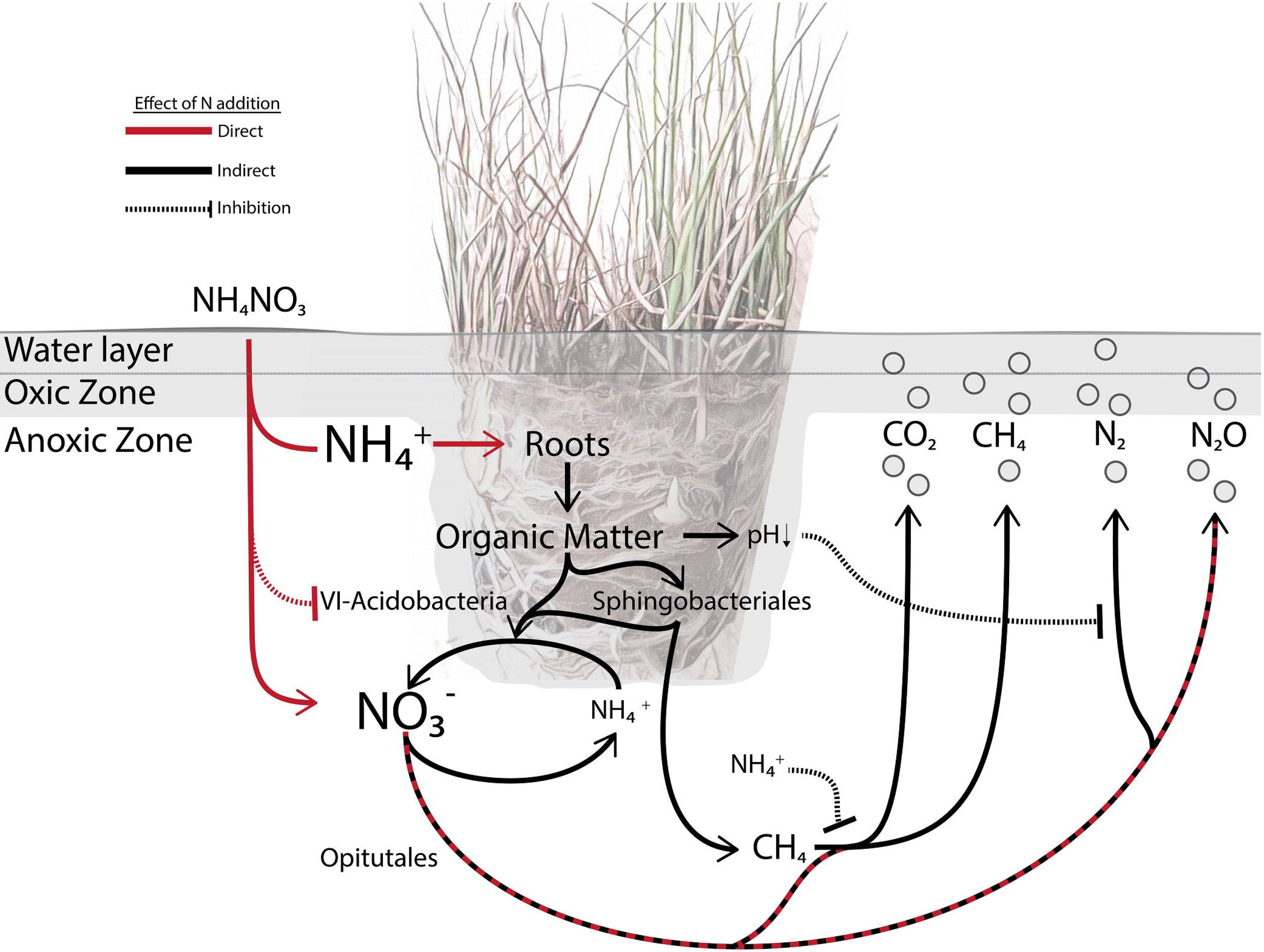
A *Juncus acutiflorus* rhizosphere microbial food web model. In the model of the *J. acutiflorus* rhizosphere, microbial processes are directly (red lines) or indirectly (black lines) influences by N deposition. *J. acutiflorus* preferentially takes up NH_4_^+^ which stimulates plant productivity and rhizodeposition of organic matter and oxygen (Van Diggelen et al., 2015). Released oxygen and labile organic matter contribute to soil acidification in addition to stimulating complex polymer degradation (Sphingobacteriales) and heterotrophic denitrifiers (Opitutales). The production of N_2_ can be affected by a drop in pH which influences the activity of complete denitrifiers. The Group-6 Acidobacteria are outcompeted at higher N availability. Recalcitrant organic matter degraded by Sphingobacteriales can enter the microbial food web and be fermented by fermenters that in turn provide substrates for methanogens (*mcr*). The activity of phosphonate lyases (*phn*) might also stimulate the production of methane while anaerobic methane oxidation also contributes to methane consumption. Additionally, methane consumption by aerobic methanotrophsthrough methane monoxgenases (*pmo*) could be inhibited by excess NH_4_^+^ (Dunfield and Knowles 1995).

### Greenhouse gas fluxes as a result of high N input

N availability has been shown to alter greenhouse gas emission dynamics in previous studies (Philippot *et al.*, 2009). Here we observed that greenhouse gas fluxes in *J. acutiflorus* incubations were stimulated as a response to increased N input (Figure 1). CO_2_ fixation rates were highest in *J. acutiflorous* incubations with high N input in the light conditions, likely due to increased photosynthetic activity of the plant and photosynthetic microorganisms. In the dark, the same *J. acutiflorus* incubations showed elevated CO_2_ emissions, likely due to increased plant and microbial respiration (Figure 1).

In this study, the highest CH_4_ emissions were observed in *J. acutiflorus* incubations with high N input, although with high variability (Figure 1C). Still, the elevated emission rates suggest that the *J. acutiflorus* rhizosphere could become a net source of CH_4_ under high N input. The total amount of CH_4_ released reflects the sum of CH_4_ production (methanogenesis) and consumption (methanotrophy). Methanogenesis has been linked to plant productivity, thought to be due to increased availability of labile organic carbon from photosynthate exudates (Whiting and Chanton, 1993;Aulakh *et al*, 2001). Furthermore, methanogens can be stimulated through an indirect priming mechanism. Labile organic matter from plant photosynthate can stimulate microbial activity responsible for degrading recalcitrant organic matter, which in turn makes this carbon source available to methanogens (Jenkinson *et al.*, 1985; Kotsyurbenko, 2005; Kotsyurbenko *et al*, 1993; Tveit *et al*, 2015). Alternatively, net CH_4_ emissions can be increased by inhibiting CH_4_ consumption, for instance through the competitive inhibition of the key enzyme methane monooxygenase by NH_4_^+^ (Bosse *et al.*, 1993; Conrad and Rothfuss, 1991).

The reduction of NO_x_ to N_2_ is often incomplete, resulting in the production of the greenhouse gas N_2_O. Incomplete denitrification occurs when microbial species do not utilize N_2_O as an electron acceptor either due to physiological constraints or induced by certain environmental conditions (Philippot, 2002; Wallenstein *et al.*, 2006). It has been observed that N fertilization has the largest impact on N_2_O emissions when considering all terrestrial ecosystems, with NO_3_^−^ availability being the main driver (Liu and Greaver, 2009). As denitrification is largely a microbial process, the composition of the microbial community plays an important role in the total amount of N emitted from soils. Representatives from a diverse set of phyla are known to denitrify (Philippot, 2002; Philippot *et al*, 2009) and denitrification rates are therefore considered to be robust to changes in the microbial community composition (Enwall *et al.*, 2005). Here we observed elevated N_2_O emissions in *J. acutiflorus* incubations under high N input, whereas there were negative N_2_O fluxes in the low N incubations. Interestingly, N_2_O emissions by bulk soil were not significantly influenced by the tested N regimes, indicating that *J. acutiflorus* plays a substantial role in stimulating N reducing microbial species, probably by supplying labile carbon. In addition there was an almost 10-fold shift in the release of N_2_O relative to N_2_ as a response to N input suggesting a high N input can shift the community towards partial denitrifiers in the rhizosphere, which is important given the strong greenhouse potential of N_2_O.

### Shifts in microbial community structure as a response to high N input

Associating microbial metabolisms (i.e., those resulting in greenhouse gas emission) to the structure of microbial communities and abiotic factors defined by the environment is essential to predict how the structure and function of these microbial ecosystems may adapt to future conditions. Bulk and rhizosphere soils contain diverse microbial communities with equally diverse metabolisms (Philippot *et al.*, 2013; Torsvik and Øvreås, 2002). It remains a challenge to understand the role that key groups play in these systems, and how they affect their environment.

We link the abundance of three bacterial orders to N input and greenhouse gas emissions (Figure 2; Table 1). The Verrucomicrobial Optitutales were associated with high N input and elevated N_2_O emissions. Members of this order are diversely associated with different rhizospheres, ranging from sugar cane to wetland plants (Dedysh *et al.*, 2006; van Passel *et al.*, 2011). They have been physiologically described as anaerobic polysaccharide utilizing bacteria that are capable of reducing NO_3_^−^ to NO_2_^−^ (Chin *et al.*, 2001). Apart from the O_2_ derived from the plant roots, which is quickly consumed by aerobic heterotrophs, wetland soils are waterlogged systems resulting in an anoxic environment. Assembled sequences from the metagenomes obtained in this study aligned to an Optitutales genome (CP016094.1), which encodes NO_3_^−^ and NO_2_^−^ reductases. Additionally, two of our assembled contigs contained open reading frames for the copper-containing nitrite reductase (NirK). It is likely that members of this order are utilizing plant derived organic matter as their electron donor and NO_3_^−^ as their electron acceptor (Figure 3).

The Sphingobacteriales from the phylum Bacteroidetes were also overrepresented in the high N input incubations (Figure 2; Table 1). Sphingobacteriales are understood as copiotrophic bacteria, referring to their ability to metabolize a wide array of carbon sources and being present at high abundances in soils with high carbon availability (Fierer *et al.*, 2007b; Padmanabhan *et al.*, 2003). In the current study, the majority of organic matter would originate from the plant as the sandy soil used had low organic matter content. Rhizodeposition in this case would be very important to groups such as Sphingobacteriales, not only as a carbon source but as an O_2_ source as Sphingobacteriales seem to be particularly sensitive to O_2_ availability. When tested for cellulolytic activity in oxic or anoxic environments they were exclusively active in the oxic treatment, suggesting that this group may require oxygenated environments for carbon degradation (Schellenberger *et al.*, 2009). Here, Sphingobacteriales were more abundant in high N input incubations and were associated with N_2_O fluxes and higher CO_2_ fixation rates, suggesting that they may benefit from oxygen and carbon derived from roots. In addition, multiple contigs from the soil metagenomes aligned to a Sphingobacteriales genome (CP003349.1), which encodes nitrate, nitrite, nitric oxide and nitrous oxide reductases. Three of these contigs encoded NirKs homologous to one found in a Sphingobacteriales genome (LGEL01000245.1). Considering findings from this study and the literature, we hypothesize that Sphingobacteriales within the *J. acutiflorus* rhizosphere could be facultative anaerobes benefiting from the elevated carbon input from the roots and utilizing available NO_x_ as electron acceptors (Figure 3).

G6 Acidobacteria were overrepresented in the low N input incubations and there was no significant difference in their abundance between bulk and rhizosphere soils. Unlike Opitutales and Sphingobacteriales, they were negatively correlated with N_2_O emissions (Figure 2; Table 1). While the G6 Acidobacteria group is not well studied, one genome (CP015136.1) was recently published (Huang *et al.*, 2016) and was shown to contain nitric and nitrous oxide reductases. 145 contigs of our metagenome aligned to this genome; however none of the assembled contigs encoded proteins involved in denitrification. Genomic and physiological studies of a closely related group (group 1 Acidobacteria) showed that they were anaerobic organoheterotrophs capable of utilizing NO_3_^−^ for respiration and NH_4_^+^ as an N source (Dedysh *et al.*, 2012), and other Acidobacteria have also been described as important soil carbon and N cyclers. However, many N-cycling reactions are restricted to particular clades indicating that these functions are heterogeneously represented across the Acidobacteria phylum (Kielak *et al.*, 2016; Koch *et al.*, 2008). Alternatively, Acidobacteria can utilze C derived from autotrophic microorganisms in anoxic environments (Meisinger *et al.*, 2007). They have been reported to utilize various plant and microbe-derived polysaccharides, like xylan, cellobiose and gellan (Janssen *et al.*, 2002; Koch *et al.*, 2008) and thrive in various soils and rhizospheres, including anoxic soils with low pH (Fierer *et al.*, 2007a; Pankratov and Dedysh, 2010). The cultured representatives of Acidobacteria have low growth rates and appear to be adapted to oligotrophic environments (Fierer et al. 2007; Jones et al. 2009). Thus, G6 Acidobacteria may not be competitive under high N availability by fast-growing (partial) denitrifiers. Together, the G6- Acidobacteria may be involved in anaerobic degradation of organic carbon from autotrophic bacteria or plant biomass, and increased N availability might reduce this group’s abundance (Figure 3).

### A model microbial food web within bulk soil and the *J. acutiflorus* rhizosphere

Increased N input poses a distinct threat to wetland ecosystems, contributing to the degradation of biodiversity and altering greenhouse gas emissions (Bobbink *et al.*, 1998; Philippot *et al.*, 2009). Plants, such as *J. acutiflorus*, influence the abundance and composition of micro-organisms living in the rhizosphere by exuding organic matter and releasing oxygen from their roots (Reinhold-Hurek *et al.*, 2015). In the current study, N addition resulted in increased productivity of *J. acutiflorus*, stimulating the effect of the plant on the microbial community but also directly affecting microbial metabolism. Based on our observations and published knowledge, we built a model of the *J. acutiflorus* microbial food web indicating how N input impacts the soil microbial community (Figure 3).

N fertilization can directly influence the soil microbial community by providing excess NH_4_^+^ and NO_3_^−^. Previous studies have shown that *J. acutiflorus* prefers NH_4_^+^ over NO_3_^−^ as N source, leading to a surplus of NO_3_^−^ in the rhizosphere (Supplementary Figure 6; van Diggelen et al. 2016). This alters N cycling dynamics, favoring microbial species capable of rapidly reducing NO_3_^−^ to N_2_O rather than to N_2_. While complete denitrification supports higher growth yields, it also is energetically more costly and thus unfavorable under lower nutrient availability (i.e., K strategy life style). The combined effect of enhanced plant derived carbon input and higher N availability stimulates heterotrophic activity, resulting in increased N_2_O and CO_2_ emissions (Figure 3). While excess NO_3_^−^ spurs anaerobic respiration, increased NH_4_^+^ concentrations can lead to an inhibition of methane oxidation, possibly contributing to the heterogeneity observed in CH_4_ emissions (Figure 1C). High N availability can also have an indirect effect by influencing plant physiology. The observed increased rates of carbon fixation by *J. acutiflorus* under high N input may result in augmented release of organic matter (including organic acids) and oxygen from the roots. This acidifies the rhizosphere soil, which can alter the activity of *nosZ* containing microbes (Liu *et al*, 2014). Additionally, elevated oxygen availability stimulates heterotrophic activity in an otherwise anoxic environment, leading to higher CO_2_ emissions. Thus, altered N input in the *J. acutiflorus* rhizosphere leads to increased greenhouse gas fluxes directly by altering the abundance of N-cycling species and indirectly through the stimulation of plant primary productivity (Figure 3).

## CONCLUSIONS

With continued anthropogenic inputs of nitrogen into wetlands, it is critical to mechanistically understand how this activity may affect globally relevant carbon and nitrogen cycling within wetlands. The results here support that under high N input, greenhouse gas emissions from the *J. acutiflorus* rhizosphere increase, shifting the system from a greenhouse gas sink to a source. Three bacterial orders, the Opitutales G6-Acidobacteria and Sphingobacteriales, respond to increased N availability and genomic evidence supports their involvement in processes leading to changes in greenhouse gas fluxes. Our view is that understanding interactions within the rhizosphere, that result in increased greenhouse gas emissions, is essential for creating management solutions aimed to address greenhouse gas emission goals, efficient agricultural practices, and conservation efforts. To move forward in our understanding of the complex dynamics within ecosystems such as the rhizosphere, future effort needs to be made in building extensive datasets that can be used to build predictive models of how these microbial ecosystems might respond under altered environmental conditions. We propose that mechanistic models, such as our *J. acutiflorus* rhizosphere plant-microbial food web model, should be used to set the framework for building such datasets.

## Acknowledgements

We would like to thank Theo van Allen for sequencing support and Sebastian Krosse, Paul van der Ven, and General Instrumentation at Radboud University for support with elemental analysis.

Funding was provided by the European Research Council (ERC AG 339880 ECOMOM) to M.S.M.J., and the Netherlands Organisation for Scientific Research (NOW) through Gravitation Grants SIAM (024.002.002), and NESSC (024.002.001) to M.S.M.J. and VENI grant 863.14.019 to S.L.

## Author Contributions

EH, SFH, JvD, LL, MJ, CL, SL, CW designed research; EH, SFH, JvD performed research; EH, SFH, JvD analyzed data; EH, SL, CW wrote the paper; All authors reviewed and agreed with the final version of the manuscript.

## Supplemental Material

**Supplemental Table 1.**

Sample overview containing the time of sampling, N load treatment and whether or the sample was bulk or rhizosphere soil. Additionally, the number of post quality filtered reads that were produced and the number of OTUs found in each sample. Finally, greenhouse gas fluxes are reported in (μmol m^−2^ d^−1^).

**Supplemental Table 2.**

Plant average dry weight and C:N were determined in different sections of the plant including the roots, shoots and rhizomes. Biomass weight was determined as dry weight. The mean values from plants receiving high N and low N are reported (Mean_High and Mean_Low). The p-value is reported as a result of a t-test comparing mean values from high and low N treatments.

**Supplemental Table 3.**

Metagenome library overview including the time of sampling (Time), number of post-QC reads (Reads), average length (Avg_len) and the standard deviation in read length (Sd_len).

**Supplemental Table 4.**

Number of reads and contigs assigned to one of three publicly available soil bacterial genomes. Number of reads each metagenome contained are reported as well as the number of contigs that were assembled that aligned to these genomes.

**Supplemental Table 5.**

Gene abbreviations.

## Supplemental Figures

**Supplemental Figure 1.**

Experimental design schema depicting sample replicates per treatment in either rhizosphere/bulk soil. Additionally, the sampling points are denoted by colored boxes.

**Supplemental Figure 2.**

The Shannon diversity index (H’) was calculated for all microbial communities. The Shannon diversity of all samples was compared from T_m_ and T_f_ (A). Diversity of experimental groups (High/Low N + Rhizosphere/Bulk) of all T_m_ (B) and T_f_ (C) samples were compared using multiple comparisons

**Supplemental Figure 3.**

Rarefaction curves with number of species observed as a function of sequencing effort (sample depth).

**Supplemental Figure 4.**

Microbial community variation was estimated by calculating the Bray-Curtis dissimilarity for each sample in a pairwise fashion resulting in a square distance matrix. These pairwise distances were then reduced to two dimensions using multidimensional scaling. A centroid was calculated for each group being compared and each sample’s Euclidean distance to its respective group centroid was calculated. T_m_ and T_f_ were compared in panel A, T_m_ (B) and T_f_ (C) samples were compared within respective groups (High/Low N + Rhizosphere/Bulk)

**Supplemental Figure 5.**

Denitrification potential from soil slurries. N and N_2_O production rates were estimated to determine potential denitrification of the soil and rhizosphere microbial communities.

**Supplemental Figure 6.**

Pore water inorganic nutrients, pH and alkalinity. Concentration of inorganic nutrients, pH and alkalinity in pore water sampled throughout the incubation.

## References

Abou SeadaMNI, OttowJCG. (1985). Effect of increasing oxygen concentration on total denitrification and nitrous oxide release from soil by different bacteria. Biol Fertil Soils 1: 31–38.

Al-GhalithGA, MontassierE, WardHN, KnightsD. (2016). NINJA-OPS: Fast accurate marker gene alignment using concatenated ribosomes. PLoS Comput Biol 12: e1004658.

ArmstrongW. (1971). Radial oxygen losses from intact rice roots as affected by distance from the apex, respiration and waterlogging. Physiol Plant 25: 192– 197.

AulakhMS, WassmannR, BuenoC, RennenbergH. (2001). Impact of root exudates of different cultivars and plant development stages of rice (Oryza sativa L.) on methane production in a paddy soil. Plant Soil 230: 77– 86.

BankevichA, NurkS, AntipovD, GurevichAA, DvorkinM, KulikovAS, et al. (2012). SPAdes: A new genome assembly algorithm and Its applications to single-cell sequencing. J Comput Biol 19: 455–477.

BardgettRD, van der Putten WH. (2014). Belowground biodiversity and ecosystem functioning. Nature 515: 505–511.

BobbinkR, HornungM, RoelofsJGM. (1998). The effects of air-borne nitrogen pollutants on species diversity in natural and semi-natural European vegetation. J Ecol 86:717–738.

BodelierPLE, LaanbroekHJ. (2004). Nitrogen as a regulatory factor of methane oxidation in soils and sediments. FEMSM icrobiol Ecol 47.

BosseU, FrenzelP, ConradR. (1993). Inhibition of methane oxidation by ammonium in the surface layer of a littoral sediment. FEMS Microbiol Ecol 13.

BragazzaL, FreemanC, JonesT, RydinH, LimpensJ, FennerN, et al. (2006). Atmospheric nitrogen deposition promotes carbon loss from peat bogs. Proc Natl Acad Sci U S A 103: 19386–19389.

BrenzingerK, DörschP, BrakerG. (2015). PH-driven shifts in overall and transcriptionally active denitrifiers control gaseous product stoichiometry in growth experiments with extracted bacteria from soil. Front Microbiol 6: 961 .

BrittoDT, KronzuckerHJ. (2002). NH_4_^+^ toxicity in higher plants: a critical review. J Plant Physiol 159: 567–584.

BuchfinkB, XieC, HusonDH. (2014). Fast and sensitive protein alignment using DIAMOND. Nat Methods 12: 59–60.

CaporasoJ, KuczynskiJ, StombaughJ. (2010). QIIME allows analysis of high-throughput community sequencing data. Nat Methods 7: 335–336.

ChinKJ, LiesackW, JanssenPH. (2001). Opitutus terrae gen. nov., sp. nov., to accommodate novelstrains of the division ‘Verrucomicrobia’ isolated from rice paddy soil. Int J Syst Evol Microbiol 51:1965– 1968.

ConradR, RothfussF. (1991). Methane oxidation in the soil surface layer of a flooded rice field and the effect of ammonium. Biol Fertil Soils 12: 28–32.

CurlEA, HarperJD. (1990). Fauna-microflora interactions. In:The rhizosphere. pp 369– 388.

DedyshSN, KulichevskayaIS, SerkebaevaYM, MityaevaMA, SorokinV V., SuzinaNE, et al.(2012). Bryocella elongata gen. nov., sp. nov., a member of subdivision of the Acidobacteria isolated from a methanotrophic enrichment culture, and emended description of Edaphobacter aggregans Koch. et al. 2008. Int J Syst Evol Microbiol 62: 654–664.

DedyshSN, PankratovTA, BelovaSE, KulichevskayaIS, LiesackW. (2006). Phylogenetic analysis and in situ identification of bacteria community composition in an acidic Sphagnum peat bog. Appl Environ Microbiol 72: 2110– 2117.

van DiggelenJMH, SmoldersAJP, VisserEJW, HicksS, RoelofsJGM, LamersLPM. (2016). Differential responses of two wetland graminoids to high ammonium at different pH values Hawkesford M (ed). Plant Biol 18: 307–315.

DunfieldP, KnowlesR. (1995). Kinetics of inhibition of methane oxidation by nitrate, nitrite, and ammonium in a humisol. Appl Environ Microbiol 61: 3129–3135.

EnwallK, PhilippotL, HallinS. (2005). Activity and composition of the denitrifying bacterial community respond differently to long-term fertilization. Appl Environ Microbiol71: 8335–8343.

FiererN, BradfordMA, JacksonRB. (2007a). Toward an ecological classification of soil bacteria. Ecology 88: 1354–1364.

FiererN, BreitbartM, NultonJ, SalamonP, LozuponeC, JonesR, et al. (2007b). Metagenomic and small-subunit rRNA analyses reveal the genetic diversity of bacteria, archaea, fungi, and viruses in soil. Appl Environ Microbiol 73: 7059–7066.

FiererN, LauberCL, RamirezKS, ZaneveldJ, BradfordMA, KnightR. (2012). Comparative metagenomic, phylogenetic and physiological analyses of soil microbial communities across nitrogen gradients. ISME J6: 1007–1017.

GallowayJN, TownsendAR, ErismanJW, BekundaM, CaiZ, FreneyJR, et al. (2008). Transformation of the Nitrogen Cycle: Recent Trends, Questions, and Potential Solutions. Science(80-) 320: 889–892.

HaicharF el Z, MarolC, BergeO, Rangel-CastroJI, ProsserJI, BalesdentJ, et al. (2008). Plant host habitat and root exudates shape soil bacterial community structure. ISME J2: 1221–1230.

Van den HeuvelRN, BakkerSE, JettenMSM, HeftingMM. (2011). Decreased N_2_O reduction by low soil pH causes high N_2_O emissions in a riparian ecosystem. Geobiology 9: 294–300.

HinsingerP, BengoughAG, VetterleinD, YoungIM. (2009). Rhizosphere: biophysics, biogeochemistry and ecological relevance. Plant Soil321: 117–152.

HoaglandDR, ArnonDI. (1950). The water-culture method for growing plants without soil. Calif Agric Exp Stn Circ 347: 1–32.

HuangS, VieiraS, BunkB, RiedelT, SpröerC, OvermannJ. (2016). First Complete Genome Sequence of a Subdivision Acidobacterium Strain. Genome Announc 4: e00469–16.

JanssenPH, YatesPS, GrintonBE, TaylorPM, SaitM. (2002). Improved culturability of soil bacteria and isolation in pure culture of novel members of the divisions Acidobacteria, Actinobacteria, Proteobacteria, and Verrucomicrobia. Appl Environ Microbiol68: 2391–2396.

JenkinsonDS, FoxRH, RaynerJH. (1985). Interactions between fertilizer nitrogen and soil nitrogen—the so-called ‘priming’ effect. J Soil Sci 36: 425–444.

JonesDL. (1998). Organic acids in the rhizosphere - a critical review. Plant Soil 205: 25–44.

JonesRT, RobesonMS, LauberCL, HamadyM, KnightR, FiererN. (2009). A comprehensive survey of soil acidobacterial diversity using pyrosequencing and clone library analyses. ISME J 3: 442–453.

KamilovaF, Kravchenko L V., ShaposhnikovAI, AzarovaT, MakarovaN, LugtenbergB. (2006). Organic acids, sugars, and l -Tryptophane in exudates of vegetables growing on stonewool and their effects on activities of rhizosphere bacteria. Mol Plant-Microbe Interact 19: 250–256.

KielakAM, BarretoCC, KowalchukGA, van VeenJA, KuramaeEE. (2016). The ecology of Acidobacteria: moving beyond genes and genomes. Front Microbiol 7: 744.

KingGM, SchnellS. (1998). Effects of ammonium and non-ammonium salt additions on methane oxidation by Methylosinus trichosporium OB3b and maine forest soils. Appl Environ Microbiol 64: 253–257.

KochIH, GichF, DunfieldPF, OvermannJ. (2008). Edaphobacter modestus gen. nov., sp. nov., and Edaphobacter aggregans sp. nov., acidobacteria isolated from alpine and forest soils. Int J Syst Evol Microbiol 58: 1114–1122.

KotsyurbenkoOR. (2005). Trophic interactions in the methanogenic microbial community of low-temperature terrestrial ecosystems. FEMS Microbiol Ecol 53: 3–13.

KotsyurbenkoOR, NozhevnikovaAN, Zavarzin GA. (1993). Methanogenic degradation of organic matter by anaerobic bacteria at low temperature. Chemosphere 27: 1745–1761.

LamersLPM, van DiggelenJMH, Op den CampHJM, VisserEJW, LucassenECHET, VileMA, et al. (2012). Microbial transformations of nitrogen, sulfur, and iron dictate vegetation composition in wetlands: a review. Front Microbiol 3: 156.

LeffJW, JonesSE, ProberSM, BarberánA, BorerET, FirnJL, etal. (2015). Consistent responses of soil microbial communities to elevated nutrient inputs in grasslands across the globe. Proc Natl Acad Sci U S A 112: 10967–10972.

LiawA, WienerM. (2002). Classification and regression by randomForest. R news 2: 18–22.

LiuB, FrostegardÅ, BákkenLR. (2014). Impaired reduction of N_2_O to N_2_ in acid soils is due to a posttranscriptional interference with the expression of nosZ. MBio 5: e01383–14.

LiuL, GreaverTL. (2009). A review of nitrogen enrichment effects on three biogenic GHGs: the CO_2_ sink may be largely offset by stimulated N_2_O and CH_4_ emission. Ecol Lett 12: 1103–1117.

LükeC, SpethDR, KoxMAR, VillanuevaL, JettenMSM. (2016). Metagenomic analysis of nitrogen and methane cycling in the Arabian Sea oxygen minimum zone. PeerJ 4: e1924.

MarschnerH, RomheldV, CakmakI. (1987). Root-induced changes of nutrient availability in the rhizosphere. J Plant Nutr 10: 1175–1184.

MeisingerDB, ZimmermannJ, LudwigW, SchleiferK-H, WannerG, SchmidM, et al. (2007).In situ detection of novel Acidobacteria in microbial mats from a chemolithoautotrophically based cave ecosystem (Lower Kane Cave, WY, USA). Environ Microbiol9: 1523–1534.

NahlikAM, FennessyMS. (2016). Carbon storage in US wetlands. Nat Commun 7: 13835.

OksanenJ, BlanchetFG, KindtR. (2015). Vegan Package.

PadmanabhanP, PadmanabhanS, DeRitoC, GrayA, GannonD, SnapeJR, et al. (2003). Respiration of 13C-labeled substrates added to soil in the field and subsequent 16S rRNA gene analysis of 13C- labeled soil DNA. Appl Environ Microbiol 69: 1614:1622.

PankratovTA, DedyshSN. (2010). Granulicella paludicola gen. nov., sp. nov., Granulicellapectinivorans sp. nov., Granulicella aggregans sp. nov. and Granulicella rosea sp. nov., acidophilic, polymer-degrading acidobacteria from Sphagnum peat bogs. Int J Syst Evol Microbiol 60: 2951:2959.

van PasselMWJ, KantR, PalvaA, CopelandA, LucasS, LapidusA, et al. (2011). Genome sequence of the verrucomicrobium Opitutus terrae PB 90-1, an abundant inhabitant of rice paddy soil ecosystems. J Bacteriol 193: 2367–2368.

PetersenW, BöttgerM. (1991). Contribution of organic acids to the acidification of the rhizosphere of maize seedlings. Plant Soil 132: 159–163.

PhilippotL. (2002). Denitrifying genes in bacterial and Archaeal genomes. Biochim Biophys Acta -Gene Struct Expr 1577: 355–376.

PhilippotL, HallinS, BorjessonG, BaggsEM. (2009). Biochemical cycling in the rhizosphere having an impact on global change. Plant Soil 321: 61–81.

PhilippotL, RaaijmakersJM, LemanceauP, van der PuttenWH. (2013). Going back to the roots: the microbial ecology of the rhizosphere. Nat Rev Microbiol 11: 789–799.

QuastC, PruesseE, YilmazP, GerkenJ, SchweerT, YarzaP, et al. (2013). The SILVA ribosomal RNA gene database proj ect: improved data processing and web-based tools. Nucleic Acids Res 41:D590–D596.

R Core Team. (2016). R: A language and environment for statistical computing. R Found Stat Comput Vienna, Austria. e-pub ahead of print, doi:10.1038/sj.hdy.6800737.

RamirezKS, CraineJM, FiererN. (2012). Consistent effects of nitrogen amendments on soil microbial communities and processes across biomes. Glob Chang Biol 18: 1918–1927.

RamsayJO, HeckmanN, SilvermanBW. (1997). Spline smoothing with model-based penalties. BehavRes Methods, Instruments, Comput 29: 99–106.

ReichPB, HobbieSE, LeeT, EllsworthDS, WestJB, TilmanD, et al. (2006). Nitrogen limitation constrains sustainability of ecosystem response to CO2. Nature 440: 922–925.

Reinhold-Hurek B, BungerW, BörbanoCS, SabaleM, HurekT. (2015). Roots Shaping Their Microbiome: Global Hotspots for Microbial Activity. Annu Rev Phytopathol 53: 403–424.

RobinsonD, HodgeA, FitterA. (2003). Constraints on the form and function of root systems. In: Springer Berlin Heidelberg, pp 1–31.

SchellenbergerS, KolbS, DrakeHL. (2009). Metabolic responses of novel cellulolytic and saccharolytic agricultural soil Bacteria to oxygen. Environ Microbiol 12: 845–861.

SchmiederR, EdwardsR. (2011). Quality control and preprocessing of metagenomic datasets. Bioinformatics 27: 863–4.

SmithCJ, DelauneRD. (1984). Influence of the rhizosphere of Spartina alterniflora Loisel. On nitrogen loss from a Louisiana Gulf Coast salt marsh. Environ Exp Bot 24: 91–93.

SoonsMB, HeftingMM, DorlandE, LamersLPM, VersteegC, BobbinkR. (2016). Nitrogen effects on plant species richness in herbaceous communities are more widespread and stronger than those of phosphorus. Biol Conserv. e-pub ahead of print, doi: 10.1016/j.biocon.2016.12.006.

TorsvikV, ØvreåsL. (2002). Microbial diversity and function in soil: from genes to ecosystems. Curr Opin Microbiol 5: 240–245.

TveitAT, UrichT, FrenzelP, SvenningMM. (2015). Metabolic and trophic interactions modulate methane production by Arctic peat microbiota in response to warming. Proc Natl Acad Sci US A 112: E2507–2516.

VerhoevenJ.T., Arheimer, B., Yin, C., & Hefting, M. M. (2006). Regional and global concerns over wetlands and water quality. Trends in ecology and evolution, 21(2), 96–103.

WallensteinMD, MyroldDD, FirestoneM, VoytekM. (2006). Environmental controls on denitrifying communities and denitrification rates: Insights from molecular methods. Ecol Appl 16: 2143–2152.

WhitingGJ, ChantonJP. (1993). Primary production control of methane emission from wetlands. Nature 364: 794–795.

ZedlerJB, KercherS. (2005). Wetland resources: status, trends, ecosystem services, and restorability. Annu Rev Environ Resour 30: 39–74.

